# Determinants of invasive species policy: print media and agriculture determine United States invasive wild pig policy

**DOI:** 10.1101/364372

**Authors:** Ryan S. Miller, Susan M. Opp, Colleen T. Webb

**Author notes:** **Correspondence:** Ryan S. Miller, 2150 Centre Avenue, Bldg B, Fort Collins, Colorado, 80526 USA, office 1.970.215.2055 cell.

## Abstract

Conflicts between wildlife, invasive species, and agricultural producers are increasing. Although direct management actions taken to mitigate these conflicts remain controversial, most stakeholders agree that better policies are needed to balance socio-economic considerations with invasive species management, wildlife conservation, and agriculture. However the interaction between societal and biological drivers that influence human-invasive species-wildlife conflict mitigation policy is poorly understood. We identify factors influencing policy leading to the establishment of a new federal program to control invasive wild pigs in the United States. We fit generalized linear models relating frequency of congressional policy activity, such as congressional hearings and reports, to frequency of print newspaper media and percent of the U.S. agricultural industry co-occurring with invasive wild pigs for 29 years preceding the establishment of the federal program in 2013. Our models explained 89% of the deviance in congressional policy activity indicating a strong linkage between congressional invasive wild pig policy activity and predictors representing the number of negative of newspaper articles, geographic distribution of print media, and percent of agricultural producers co-occurring with invasive wild pigs. These effects translated to 3.7% increase in the number of congressional policy actions for every additional five states with negative news media. Invasive wild pig congressional policy activity increased 6.7% for every additional 10 negative newspaper articles. Increases in co-occurrence of agriculture and invasive wild pigs had the largest effect, for every 1% increase in co-occurrence there was a 41% increase in congressional policy activity. Invasive wild pig congressional policy activity that explicitly addressed livestock increased at nearly twice the rate of policy activity addressing crop agriculture. These results suggest that agriculture and media coverage may act as determinants for invasive species policy. Our approach may provide early insight into emerging policy areas enabling proactive policy development by agencies or early engagement by scientists to find solutions before the policy area becomes grid locked. Our results can also support policy and program evaluation providing a means of determining if the implemented policies match the original policy determinants ensuring best alignment with public, environmental, and stakeholder interests.

## Introduction

Conflicts between wildlife, invasive species, and agricultural producers are increasingly challenging management agencies (Krebs et al. 1998, Miller et al. 2013, Miller and Sweeney 2013). Policy to address human-wildlife-invasive species conflicts is often controversial (Messmer 2009, Crowley et al. 2017). Although policies to manage interactions among invasive species, wildlife, and agriculture has been identified as critically important (Jones et al. 2013, Paini et al. 2016) and there have been focal studies (McBeth and Shanahan 2004, Kokotovich and Andow 2017), little research has been conducted to identify societal factors that stimulate federal policy development to address these conflicts. This gap is particularly evident for invasive species conflicts which can have complex societal and management drivers (Estévez et al. 2015, Crowley et al. 2017). The drivers that often influence policy development to address social problems include problem severity, interest group involvement, media coverage, and public perceptions (Gilliam Jr and Iyengar 2000, Soroka 2003, Walgrave et al. 2008, Baumgartner and Jones 2010).

Agriculture and wildlife policy development is often exacerbated by invasive or exotic animals (Pimental 2007). North America and in particular the United States (U.S.) has the greatest number of non-native invasive species globally, causing an estimated $46 billion in damage annually (Pimental 2007, Turbelin et al. 2017). The invasive wild pig (IWP) *Sus scrofa*, often referred to as feral hog, feral pig, feral swine or wild boar, are the most abundant free-ranging, exotic ungulate in the U.S. and are the descendants of Eurasian Russian boar (*Sus scrofa linnaeus*), feral domestic swine (*Sus scrofa domestica*), and hybrids between the two (Mayer and Brisbin 1991, Keiter et al. 2016). Since the 1960s IWPs have expanded their range to at least 38 states and 3 provinces in Canada impacting ecosystems, wildlife, and agricultural (Bevins et al. 2014, Brook and Beest 2014, Michel et al. 2017, Miller et al. 2017). Environmental and agricultural damage caused by IWPs is at least USD$1.5 billion annually (Pimental 2007, Bevins et al. 2014, Anderson et al. 2016). The rooting behavior and omnivorous diet of IWPs can have ecosystem-level effects on native plant and wildlife communities and has complicated threatened and endangered species conservation with at least 87% of imperiled species potentially impacted in the United States (Barrios-Garcia and Ballari 2012, McClure et al. 2018).

There is a diversity of public attitudes toward IWPs depending on if the animals are seen as pests, disease hazard, commodity, source of income, or recreational resource (Tisdell 1982, Izac and O'Brien 1991, Caplenor et al. 2017). Economic impacts and perceived problem severity often differ regionally. Despite economic costs IWPs are also valued as an economically important hunting resource with 35 states currently allowing public hunting and in 11 states IWPs are under the jurisdiction of the state game and fish agency (Centner and Shuman 2015, Group' 2016, Caplenor et al. 2017). Izac and O'Brien (1991) found that these perceptions changed with location, overtime and how an individual may be affected. This diversity of public opinion influences public discourse concerning how IPWs should be managed subsequently affecting institutional (e.g. governmental agencies) problem identification and implementation of policies to control or mitigate the damage caused by IWPs.

Conflict over public policies can be decomposed into the scope of participation by affected parties and how the problem and its solutions are perceived (Schattschneider 1960). These two aspects of policy conflict are useful in understanding the relative contribution of interest groups and public perception of the problem to policy development. Extensive work in the policy analysis literature indicates that identifying the visible, and presumably most affected (often economically affected), participants in a policy issue is central to understanding why some policy problems receive attention by governmental institutions resulting in policy development (Kingdon and Thurber 1984, Baumgartner and Jones 1991, Jones and Baumgartner 2004, Baumgartner et al. 2009, Baumgartner and Jones 2010). This is often tied to increasing problem severity that often results in increased lobbying by interest groups and can be a significant stimulus for the adoption of policy innovations (Sapat 2004). The dominant conceptualization of a policy problem and the solutions, often referred to as the ‘policy image’, is also important in understanding policy development (Barrilleaux et al. 2017). Media coverage of public issues – both quantity and tone (i.e. negative or positive) - has been widely recognized as an important driver in shaping national public perception and perceived importance of policies issues thus influencing government institution policy agendas (Gilliam Jr and Iyengar 2000, Walgrave et al. 2008, Baumgartner and Jones 2010). Media coverage is generally thought to effect government policy agendas by increasing the relative salience (i.e. importance) of a particular pubic issue and increasing policy image coalescence often early in the policy process (Elder and Cobb 1983, Weart 1988, Soroka 2003, Baumgartner and Jones 2010). Salience and coalescence of a policy image is often translated into pressure on government officials to prioritize development of policy solutions.

There is mixed evidence for how these factors - media coverage, public perception, problem severity, and interest groups - may act together to influence policy generation (McCombs and Shaw 1972, Funkhouser and Shaw 1990, Entman 1993, Koch-Baumgarten and Voltmer 2010) and there is generally poor understanding of how they may influence policy addressing conflicts among invasive species, wildlife, and agricultural (McBeth and Shanahan 2004, Lodge and Matus 2014). Our objectives in this study were to characterize the relative contribution of these factors to the development of national invasive species policy. Specifically we wanted to understand 1) the significance of public policy image on congressional policy activity (e.g. congressional hearings and reports) to address invasive species conflict; 2) determine the contribution of problem severity and the resulting pressures on government institutions from interest groups to develop policy solutions; and 3) to identify predictors of policy activity for informing invasive species management and policy; specifically program assessments and new program development. To investigate these relationships we use 29 years of federal congressional policy activity leading to the establishment of the Animal Plant Health Inspection Service (APHIS) National Feral Swine Damage Management Program in 2013 (federal government fiscal year 2014) (USDA 2013). The broader goal of this analysis is to provide a mechanistic understanding of factors contributing to invasive species policy enabling improved development of policies to manage conflicts with invasive species.

## Methods

### Congressional policy activity data

A systematic search of the Federal Digital System (FDsys) maintained by the United States Government Printing Office (GPO 2014) was used to generate data describing congressional policy activity related to IWPs. We use the term ‘policy activity’ in its broadest definition referring not only to operational policies of government (e.g. code of federal regulations) but also including all official discourse related to the development of policy (e.g. committee hearings and congressional reports). The FDsys is an official repository of all official publications from all branches of the United States Federal Government and currently contains over 7.4 million electronic documents from 1969 to present. Our search included congressional hearings, congressional record, congressional reports, bills, and changes to the code of federal regulations from 1985 until 2013 when the APHIS National Feral Swine Damage Management Program was established. FDsys documents included in our study contained any of the following search terms: ‘feral swine’, ‘feral hog’, or ‘feral pig’, ‘wild swine’, ‘wild hog’, or ‘wild pig’.

Each document was considered an independent policy action, and the number of documents by year was tallied to generate count data by document type and the primary agricultural commodity (livestock or crop) the document addressed. Our method may have included documents which were not specifically addressing IWP related policy. Because it was prohibitive to manually assess 476,000 pages of policy documents to evaluate inclusion error, we randomly sampled 5% of the documents to determine if they addressed IWP policy allowing inclusion error greater than 4.5% with 95% certainty to be detected (Valliant et al. 2013). Based on the results of this assessment we assumed that if the document contained a reference to IWPs the issue was either on the policy agenda or influencing the agenda in some way (Baumgartner and Jones 2010).

The policy process is an often nonlinear multi-stage cycle that can be characterized into at least six primary stages that include, 1) issue emergence and problem formation, 2) agenda setting, 3) formulation 4) policy adoption, 5) implementation, 6) evaluation (Brewer and DeLeon 1983, Anderson 1984). These six stages often overlap, often have additional hierarchy of stages, and are not always required for policy to be in-acted (Sabatier and Weible 2014). To support our statistical analyses the policy documents were used to identify the primary policy stages that IWP policy likely experienced. We used the policy stage heuristic to determine the policy stages that IWP policy likely experienced (Brewer and DeLeon 1983, Anderson 1984) (see Appendix S1). These stages were then used in post-hoc analysis of the contribution of our policy model predictors to overall policy activity (see Policy models methods).

### Print media data

Data on media reporting of IWP related topics was generated using a search of four major news consolidators – Newsbank, LexisNexis, EBSCO Information Services, and ProQuest (EBSCO 2016, LexisNexis 2016, NewsBank 2016, ProQuest 2016). We used similar search criteria to that used for the policy activity data. Our review was restricted to print newspaper articles (here after referred to as articles) published from 1985 to 2013 in the United States contained in these databases. In order for an article to be included it must have contained the terms ‘feral swine’, ‘feral hog’, or ‘feral pig’, ‘wild swine’, ‘wild hog’, or ‘wild pig’ in the title or lead into the article. Articles published by the same newspaper and author on the same date were considered duplicates and removed. The data were tallied by year to generate three annual predictors, the number of articles, the number of different newspapers, and the number of states with at least one article. A state was considered to have an article if the newspaper was located in the state. For national newspapers such as the New York Times, the head office was used to determine the state (e.g. the New York Times would be located in New York).

To generate a measure of article tone, each article headline was classified as positive or negative (see detailed methods in Appendix S2). Our assumption was that the article headline summarized the content and sentiment of the article representing the overall tone of the article and has previously been used as an index of tone (Dodds et al. 2011, Dodds et al. 2015). We classified articles as having positive or negative tone using an index previously described by Rinker (2013) and Liu (2015) using the polarity function in the Qualitative Data and Quantitative Analysis package (Rinker 2013) within the R computing environment (R Core Team 2016). Briefly, the polarity algorithm uses a word sentiment (positive or negative) dictionary to identify and assign a score to polarized words in the article headline (Hu and Liu 2004). The words adjacent to the polarized word, often referred to as the context cluster, are used to weight the score for the polarized word. The weighted scores for each polarized word within the article headline are summed and divided by the square root of the word count for the headline yielding an unbounded score for article tone in which zero is perfectly neutral, positive values indicate positive tone and negative values indicate negative tone.

We were interested in the annual influence of article tone and newspaper tone on policy activity. To this end the polarity scores for each article were used to calculate the mean article tone (article tone) in each year across all articles. To calculate the annual mean newspaper tone we first calculated the mean article tone by newspaper in each year. This represented the mean article tone for each newspaper in each year (i.e. tone of articles published by each newspaper). We then calculated the mean newspaper tone across all newspapers in each year providing a measure of mean annual newspaper tone (newspaper tone). A detailed description of the methods used to generate article and newspaper tone using the polarity index are described in Appendix S2.

### Increasing problem severity and interest group pressures

The most affected participants, often economically, in a policy area often determine policy outcomes (Kingdon and Thurber 1984, Baumgartner and Jones 1991, Jones and Baumgartner 2004, Baumgartner et al. 2009, Baumgartner and Jones 2010). Increasing problem severity and economic losses often increase lobbying by interest groups and is considered a primary mechanism of interest group engagement (Sapat 2004) In the case of IWPs there are several potentially affected interest groups including wildlife conservation, sportsmen, and agriculture. Agriculture likely has the largest economic impacts resulting from IWPs (~USD$1.5billion) although the economic impacts to sportsmen and wildlife are not as easily estimated or available (Pimental 2007). We limited our investigation to agricultural interest groups which are likely the most economically affected group.

To generate a measure of potential interest group pressures on governmental institutions (here after institutional pressure) we developed a proxy variable describing the co-occurrence of agricultural producers and IWPs. Data describing the nationwide distribution (presence/absence) of wild swine at the county level was compiled from the Southeastern Cooperative Wildlife Disease Study (SCWDS) (Corn and Jordan, SCWDS 2013) and two publications Waithman et al. (1999) and Hanson and Karstad (1959). These data represent the known nationwide county level distribution of IWPs for 39 states and 1,521 counties over the past 50 years and have been used to forecast the spread of IWPs (Snow et al. 2017), estimate the probability of occurrence (McClure et al. 2015), and determine agricultural producers at risk for damage from IWPs (Miller et al. 2017). For each year and county with IWP presence the number of agricultural producers was determined using the National Agricultural Statistics Service (NASS) data (USDA 2014). Because data describing the distribution of IWPs were not available for all years and represent a sample of the known distribution of IWPs over time, models were fit to these data to estimate the total number of agricultural producers in counties where IWPs occur for each year of policy data (McClure et al. 2018). To improve estimation for earlier years where IWP distribution data is sparse we used all IWP – producer data to fit the models. We determined relative support in the data using four candidate models - linear, exponential, power, and logistic - to describe the phenomenological change in national co-occurrence of IWPs and agriculture. Akaike information criterion with a correction for small sample size (AICc) was used to determine the best approximating model. This model was used to predict the mean national proportion of agricultural producers co-occurring with IWPs for each year from 1985 to 2013 and was used as a predictor in the policy models.

### Policy models

We evaluated support for competing models portraying the relationship between the annual count of policy actions (response variable) and six variables of interest measuring annually the 1) number of articles, 2) number of newspapers with articles, 3) number of states with articles, 4) tone for articles, 5) tone for newspapers, 6) and the proportion agricultural producers in regions with IWPs, here on referred to as agriculture. Specifically, these independent variables represent three hypotheses about specific mechanisms that resulted in congressional policy activity that eventually resulted in the establishment of a national program to address the problem.

*Problem Salience:* An increase in the number of articles, newspapers with articles, and the number of states with newspaper articles would increase the salience of the policy image increasing congressional policy actions.

*Problem Coalescence:* An increase in negative newspaper article tone for IWPs represents coalescence of the policy image increasing congressional policy activity.

*Institutional Pressures:* Increasing the number of agricultural producers in IWP regions is related to increasing problem severity and results in increased pressures on Federal government institutions by interest groups (i.e. lobbying) to find a policy solution thus increasing congressional policy activity.

We used multi-model inference within an information-theoretic framework to estimate model parameters describing the probability of congressional policy actions related to IWP co-occurrence with agriculture and media tone (Burnham and Anderson 2002, Burnham et al. 2011). All models used a Poisson error structure and were fit using a generalized linear model (GLM) with a log link function having the form:

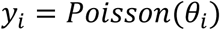

Where *y_i_* represented congressional policy actions in year *i* and *θ_i_* is the Poisson rate parameter representing the mean number of congressional policy actions in a year. The mean number of congressional policy actions, *θ_i_*, was a function of covariates on the logarithmic scale represented as:

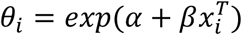

Where *α* was the intercept describing the estimated background number of congressional policy actions common across years, *β* is a vector of regression coefficients corresponding to 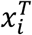, the transpose of the of the *m* × 1 vector of covariates associated with the *i*th year in the congressional policy data.

Among year correlation in the number of congressional policy actions was investigated using generalized linear autoregressive moving average (GLARMA) models (see Appendix S3), however models that accounted for serial dependence in the response variable were not significantly different than a GLM indicating between year dependence in policy actions was not important thus we did not include serial dependence in our models.

Akaike information criterion with a correction for small sample size (AICc) was used to assess the relative information content of the models. We fit all subsets of the global model and computed model-averaged regression coefficients, unconditional standard errors (SE), cumulative AICc weights of evidence as a measure of variable importance and 95% confidence intervals (Burnham and Anderson 2002, 2004, Burnham et al. 2011). Because the policy data are over-dispersed we used a shrinkage estimation approach to produce unconditional model averaged parameter estimates, in which covariates that did not appear in a particular model subset were assigned coefficients of zero to avoid biasing coefficient estimates away from zero (Burnham and Anderson 2002). Our interpretation of the explanatory power of the regression coefficients in our model was guided by: 1) the weights of evidence, ranging from 0 to 1.0, where higher weights indicated greater relative importance; 2) the 95% confidence interval for each regression coefficient that did not overlap zero; and 3) effect sizes indicated by each regression coefficient.

The final inferential model was used to estimate the mean annual contribution of each predictor (i.e. influence on the amount of policy activity in each year) to policy activity across the 29 years investigated. We also estimate the relative contribution of livestock and crop agriculture to annual federal policy activity for IWPs. Differences in the contribution among predictors to policy activity was determined using Tukey’s honest significant difference test (Kleinbaum et al. 2013) for each of the policy stages identified using the policy documents. Maximum likelihood estimates, confidence intervals on model parameters, significance statistics, and AICc values were obtained using MuMIn Multi-Model Inference package (Barton and Barton 2015) available in R (R Core Team 2016).

### Model assessment and validation

We assessed model fit using k-fold cross-validation which contrasts the number of policy actions predicted by the model and the observed frequency of policy actions (Kohavi 1995) and calculated adjusted D^2^ which is a quality-of-fit statistic (Guisan and Zimmermann 2000, Weisberg 2005). To implement k-fold cross validation we randomly divided the policy action data among four cross-validation folds using Huberty’s rule (Huberty 1994). We used all possible sets of three folds to fit the predictive model. Employing multi-model averaging, we then used the model to predict the fourth withheld fold. Results of 100 iterations of this process, each with a new random allocation of data across the folds, were averaged to avoid dependency on a single random allocation of data. We then calculated a Pearson correlation between predicted values and the observed number of policy actions. The Pearson correlation was used to assess the performance of our final model. Because validation results can be sensitive to binning method (Boyce et al. 2002), we applied and compared the results using a quantile binning method for 4, 10 and 20 bins. We also calculated adjusted D^2^ which is a measure of the amount of deviance the model accounts for adjusted by the number of observations and the number of model parameters (Guisan and Zimmermann 2000, Weisberg 2005). Adjusted D^2^ allows direct comparison among different models. Adjusted D^2^ was calculated using the modEvA package (Barbosa et al. 2016) and cross-validation was implemented using custom code in the R statistical software (R Core Team 2016).

## Results

### Congressional policy activity

Our search of FDsy for policy documents identified 421 documents related to IWPs (Figure 1). The policy documents represented three primary policy stages of increasing policy activity described by (Anderson 1984) – issue emergence and problem formation, agenda setting, policy formulation and implementation (see Appendix S1). The period from 1985 to 1993 showed no observed policy activity followed by a brief period of regulatory activity from 1994 to 1998 (e.g. changes to the federal register and the code of federal regulations) indicating that IWP policy was not yet formally present on institutional agendas and the issue was still emerging (Anderson 1984, Baumgartner and Jones 1991). From 1999 to 2006 policy activity on IWPs began in the form of congressional hearings indicating that the topic of IWPs had reached the institutional agendas of policy makers (Baumgartner and Jones 1991, 2010). The last stage was dominated by policy formulation and implementation from 2007 to 2013 which accounted for 64% of the total policy activity and comprised both regulatory and distributive policies (i.e. allocation of fiscal resources to address specific issues related to IWPs) and culminated in the establishment of a national program to address IWP damage.

**Figure 1.**
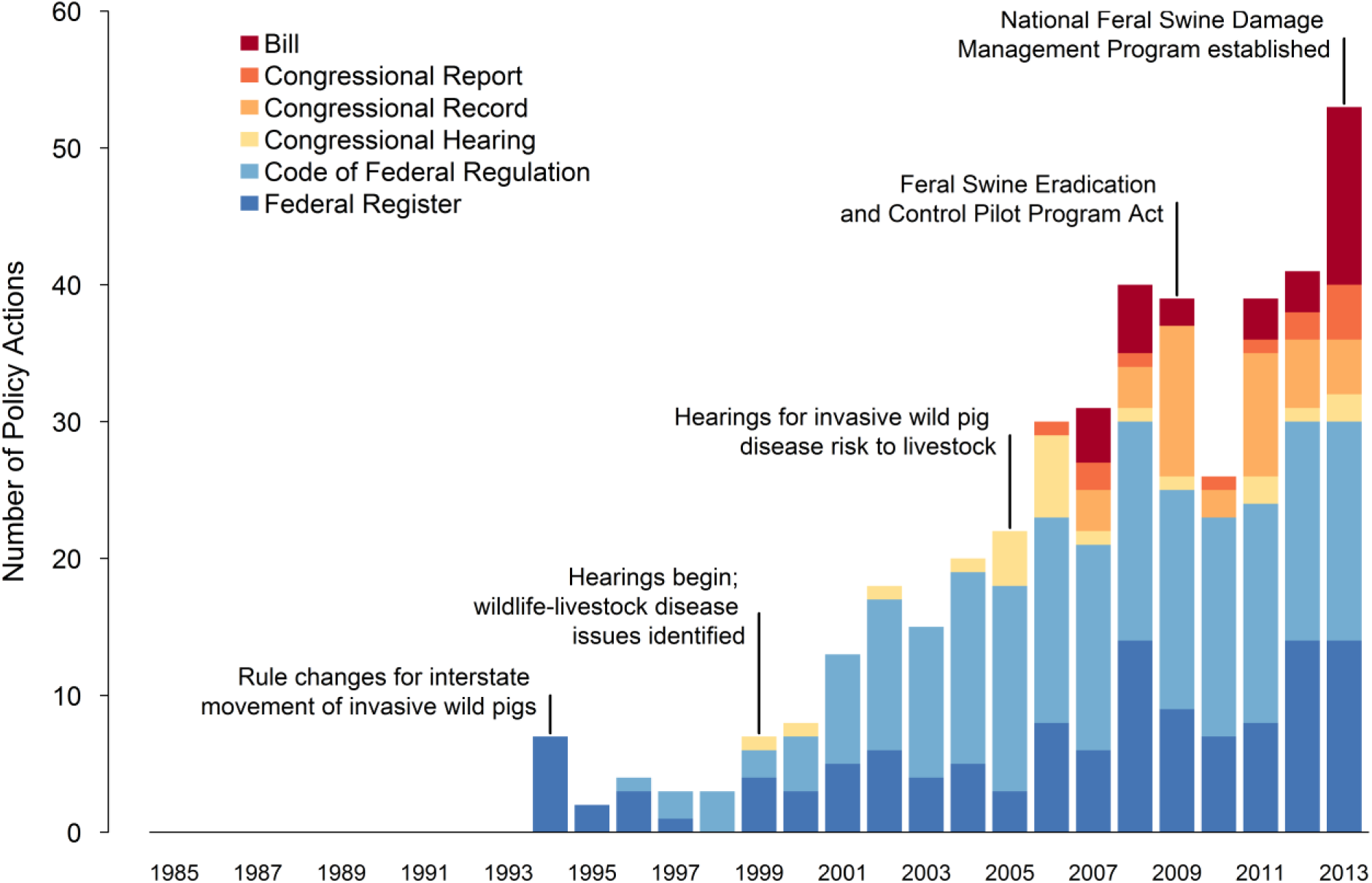
Congressional policy activity and major policy milestones for invasive wild pigs from 1985 through 2013. Blue colors represent regulatory policy activity associated with changes to the code of federal regulations and federal register. Yellow and orange colors represent the agenda setting policy stage and include activity associated with congressional hearings, reports, and record. Red indicates the policy formation and implementation stage and includes bills addressing invasive wild pig policy.

### Invasive wild pig print media

We identified 980 unique articles from 452 newspapers related to IWPs between 1985 and 2013 (Figure S2.1). The number of articles, number of newspapers and number of states with newspaper articles were relatively constant prior to 1998 with a rapid increase in articles, newspapers, and states after 1999. This period from 1999 to 2013 accounted for 96% of articles and 85% of newspapers. The number of states with wild swine related newspaper articles continued to increase throughout the study period with 47 states having at least one article.

### Co-occurrence of invasive wild pigs and agriculture

The co-occurrence of IWPs and agriculture expanded at an increasing rate from 1959 until 2013 and was best approximated by a logistic model (Table 1). For our study period the national proportion of agricultural producers in regions with IWPs increased from 0.17 in 1985 to 0.41 in 2013 (Figure 2 panel B) and is similar to changes in co-occurrence of domestic animal production and IWPs found in at least one other study (Miller et al. 2017). This represented an annual rate of increase of 1.01 (stdev <0.01) in the co-occurrence of agricultural producers and IWPs during this period. Based on the strong predictive capacity of this distribution (adjusted R^2^ = 0.99) it was used as a predictor in the policy models to represent the annual number of agricultural producers potentially impacted by IWPs and provided a surrogate variable for changes in institutional pressure potentially resulting from interest group activity (i.e. lobbying).

**Table 1.** Candidate models use to estimate the county scale co-occurrence of invasive wild pigs and agricultural producers that was used as a predictor in the policy models. The best approximating model in the candidate set of models was a logistic model and had good predictive capacity with an adjusted R^2^ = 0.99.

| Candidate Models | df | AICc | $\Delta$ AICc | Aikaike Weight | Adjusted $R^2$ |
| --- | --- | --- | --- | --- | --- |
| Logistic | 7 | -58.04 | 0.00 | $7.63 \times 10^1$ | 0.99 |
| Linear | 7 | -36.31 | 21.73 | $1.46 \times 10^5$ | 0.92 |
| Exponential | 7 | -27.74 | 30.30 | $2.01 \times 10^7$ | 0.99 |
| Power | 7 | -27.16 | 30.88 | $1.51 \times 10^7$ | 0.99 |

**Figure 2.**
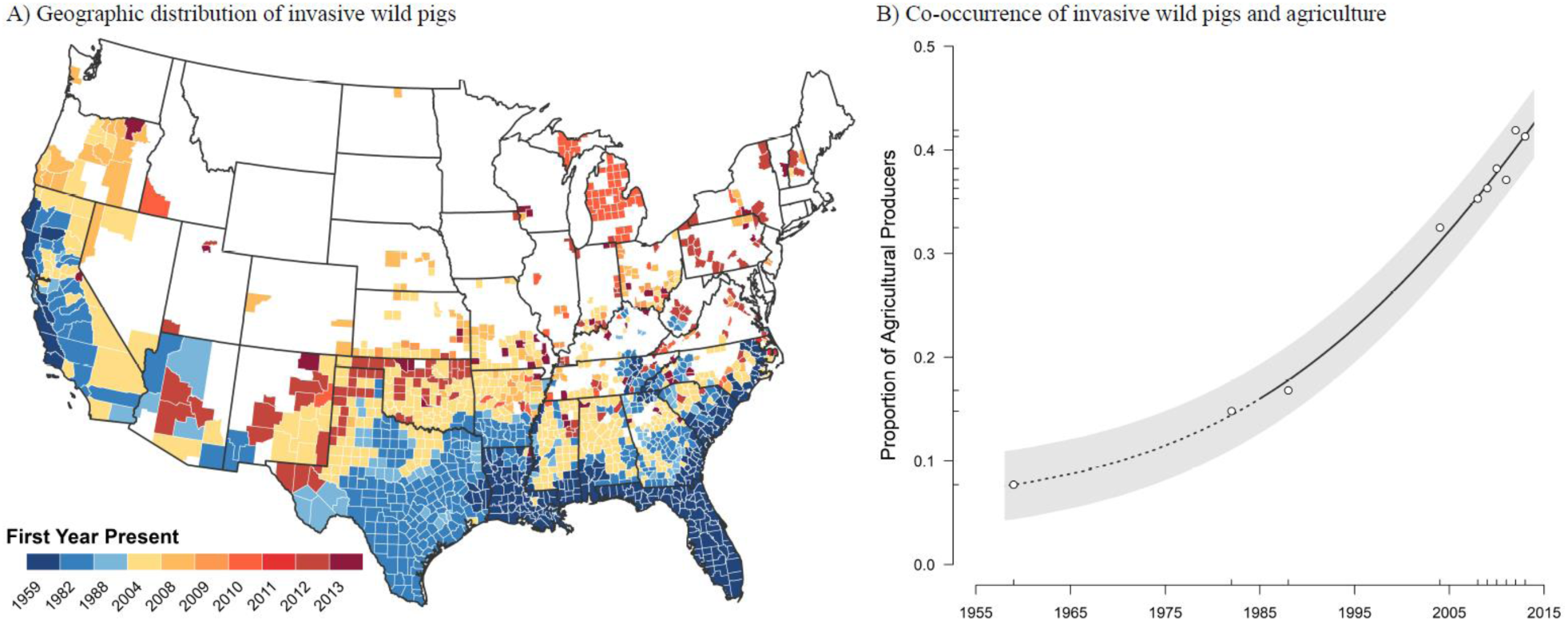
Change in distribution of invasive wild pigs and co-occurrence with agriculture in the United States. Panel A illustrate the change in the geographic distribution of invasive wild pigs in the United States. Years indicate the first year invasive wild pigs were identified within the county based on the SCWDS data. Blue scaled colors represent ‘historical’ range from 1959 until 1988. Yellow and orange scaled colors represent ‘contemporary’ range from 2004 until 2013. Panel B reports the increase in county level co-occurrence of invasive wild pigs and agriculture. Circles represent observed proportion of counties in the United States in which invasive wild pigs and agriculture co-occur. Black line (solid and dotted) denotes logistic model estimated mean and gray band is the 95% prediction interval. Solid line indicates the estimated mean used as a predictor for the years of our study – 1985 to 2013. The annual rate of increase was estimated as 1.01 (stdev <0.01) from 1959 to 2013 with the estimated inflection year being 2034 with 69.9% of agriculture co-occurring with invasive wild pigs. The model had good predictive capacity having an adjusted R^2^ = 0.99.

### Policy models

Based on the final inferential model, policy activity was most strongly associated with the number of states with newspaper articles, media source tone, newspaper article tone, and the number of agricultural producers co-occurring with IWPs (Table 2 and Table 3). Covariates representing each of these four factors had high AICc weights of evidence and 95% confidence intervals that did not include zero indicating high predictive importance. The final model accounted for most of deviance with an adjusted D^2^ of 0.89. Cross-validation indicated that the final model had strong predictive capacity. The quantile binning method produced similar Pearson correlations of 0.969 (4 bins), 0.915 (10 bins) and 0.957 (20 bins) between predicted policy actions and the observed policy actions.

**Table 2.**
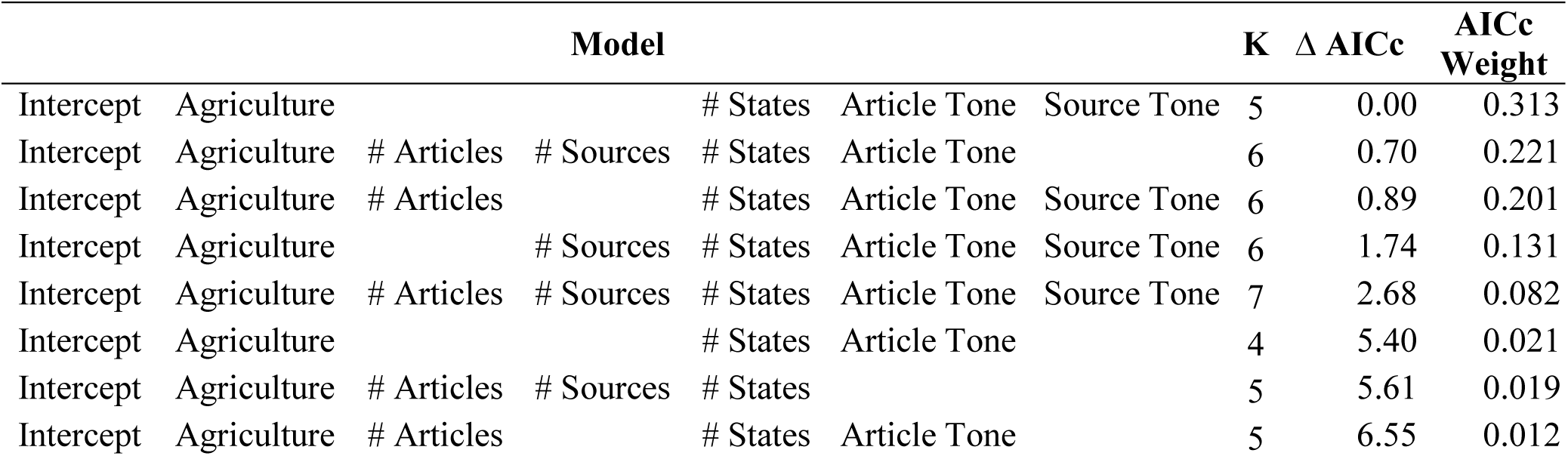
Candidate set of models used in the model averaging procedure to generate the final inferential model. These eight models account for 99.9% of the AICc weight given the candidate set of 64 models. The null model (intercept only) was ranked as the least informative model and the top model was 485 AICc units better (i.e. lower) than the null model suggesting the selected covariates approximated the data well.

**Table 3.** Model-averaged parameter estimates for the final inferential model describing the relationship between invasive wild pig policy activity, news media, and the amount of agriculture co-occurring with invasive wild pigs.

| Parameter | Odds Ratio | Estimate | Unconditional Standard Error | Unconditional Confidence Interval |  | AICc Weight |
| --- | --- | --- | --- | --- | --- | --- |
|  |  |  |  | 2.5% | 97.5% |  |
| All agriculture | 4.09 | 1.41 | 0.25 | 1.32 | 1.50 | 1 |
| Livestock agriculture | 4.08 | 1.41 | 0.34 | 1.32 | 1.49 | 1 |
| Crop agriculture | 3.43 | 1.23 | 0.53 | 0.96 | 1.51 | 0.92 |
| # States with news media | 2.08 | 0.73 | 0.17 | 0.69 | 0.78 | 1 |
| Newspaper article tone | 1.95 | 0.67 | 0.23 | 0.59 | 0.75 | 0.98 |
| Media source tone | 1.14 | 0.13 | 0.08 | 0.11 | 0.15 | 0.73 |
| # Newspaper articles | 1.98 | 0.68 | 1.04 | -0.88 | 2.25 | 0.53 |
| # Media Sources | 0.53 | -0.64 | 1.36 | -2.72 | 1.44 | 0.46 |

Parameter estimates, odds ratios, unconditional confidence intervals, and AICc weights for the predictor variables considered are presented in Table 3. The number of states with IWP related newspaper articles was a positive predictor of IWP policy activity (odds ratio = 2.08). For every additional 5 states with newspaper headlines related to IWPs there was a 3.65% increase in the number of policy actions. Increasing negative tone of the number of newspaper articles (odds ratio = 1.95) and the number of media sources (odds ratio = 1.14) increased the number of IWP policy actions. That is for every 10 negative newspaper articles and 10 negative media sources IWP policy activity increased by 6.7% and 1.3%. The number of agricultural producers in regions with IWPs was the most significant predictor of policy actions (odds ratio = 4.09); that is for every 1% increase in the proportion of agriculture in regions with IWPs policy activity increases by 41%. The amount of agriculture in wild swine regions was also a significant predictor of livestock (odds ratio = 4.08) and crop (odds ratio = 3.43) specific policy activity for IWPs. Figure 3 illustrates the functional relationship between increasing co-occurrence of agriculture and IWPs and the resulting change in IWP policy activity.

**Figure 3.**
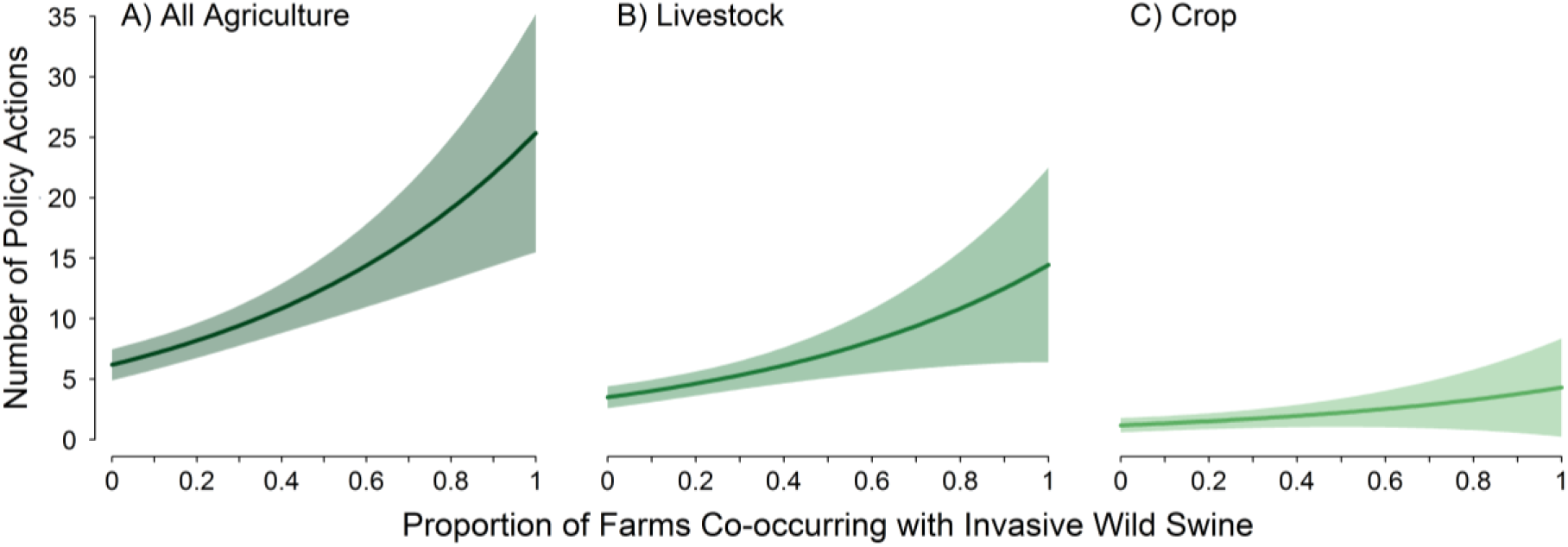
Mean functional relationship between invasive wild pig congressional policy activity and the national proportion of agriculture co-occurring invasive wild pigs in the United States. Solid black line indicates predicted mean relationship and gray band indicates unconditional 95% confidence interval. Panels represent the functional relationship for (A) all invasive wild pig policy activity, (B) invasive wild pig policy activity specific to livestock agriculture, and (C) invasive wild pig policy activity specific to crop agriculture.

The predicted contribution of the four most important predictors to policy activity changed across the 29 years evaluated and differed for the three policy stages identified (Figure 4 and Figure S1.1). During the problem formation stage from 1985–1998 all media predictors contributed greater to policy activity than agriculture (p-values < 0.001) and source tone contributed more than either article tone or states with media (p-values < 0.001) (see Appendix S4). There was no difference in the mean contribution among the predictors during the policy agenda setting stage from 1999 to 2006 (p-values > 0.05). During the policy formation and implementation stage from 2007 to 2013 agriculture contributed more (p-values < 0.05) to policy activity compared to media predictors and there was no difference in the contribution among media predictors (p-values > 0.05). Livestock specific policy activity was greater (p-values < 0.05) than crop policy activity across all years. The mean annual predicted contribution of agriculture to policy activity varied the most, with a 54.9% change from 5.5% of policy activity in 1985 to 60.7% of policy activity in 2013. Both media source and newspaper article tone had declining mean annual contribution to policy activity, declining 37.5% and 17.2%. Combined media source and article tone contributed on average 30.5% of policy activity in 2013 compared to a combined 71.7% in 1985. The number of states with newspaper articles on average contributed a consistent amount annually (22.8%) to policy activity across all years.

**Figure 4.**
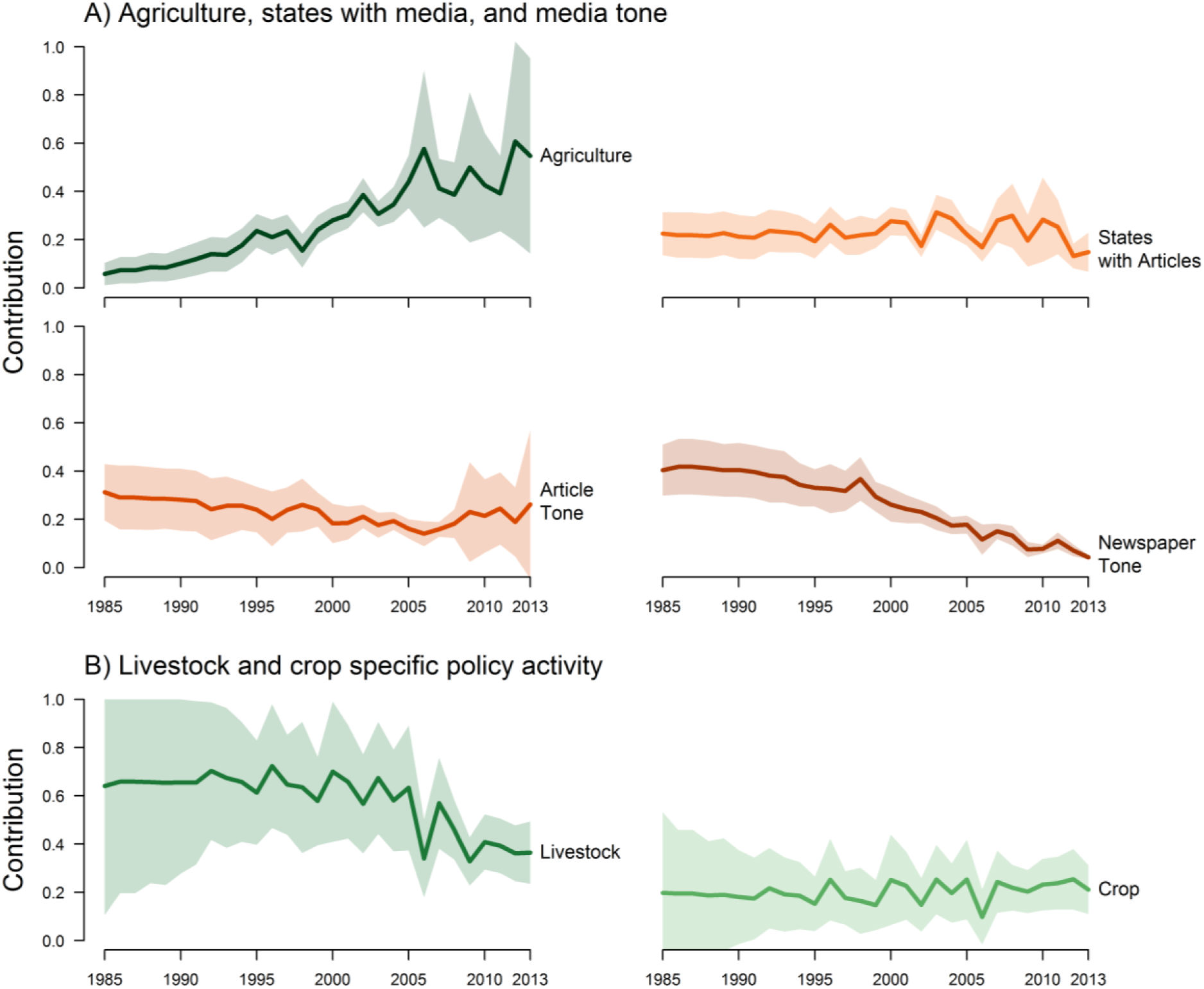
The model predicted change in relative contribution of predictors to annual policy activity for invasive wild pigs. Panel A contrasts the changes in the annual contribution of agriculture, article and newspaper tone, and number of states with newspaper articles. Panel B describes the relative contribution of livestock and crop specific activity to overall invasive wild pig policy. Solid lines indicate mean contribution and shaded region represents 95% confidence interval.

## Discussion

Our models found a linkage between policy activity and four predictors representing number of states with media, media tone and agriculture. These predictors have been described in previous studies as representing policy image salience, policy image coalescence, and institutional pressures (Elder and Cobb 1983, Kingdon and Thurber 1984, Weart 1988, Gilliam Jr and Iyengar 2000, Schnell 2001, Soroka 2003, Sapat 2004, Walgrave et al. 2008, Baumgartner and Jones 2010). We found the predicted contribution of these predictors to policy activity changed across the 29 years analyzed and differed for the three policy stages indicating the development of federal IWP policy went through a continuum of policy development. Understanding how these predictors, that serve as proxy measures of mechanisms influencing policy processes, contribute to policy development can provide a better understanding of important latent processes that give rise to national policies. This in turn can support the development of programs and policies that best address the social issues underlying these problems.

The emergence of invasive IWPs as a policy issue was characterized by decades of general inattention and no observed policy activity (Figure 1; Appendix S1). Media predictors contributed most to policy activity during the first stage of policy activity and may have acted to increase policy image salience and coalescence (Figure S1.1). Our results suggest that for IWP policy, increasingly negative news media may have acted as a mechanism for influencing initial policy activity. Previous studies have proposed that increasing news media, specifically negative news media, indicates increasing public policy image coalescence and policy issue salience (Jones and Baumgartner 2004, Baumgartner and Jones 2010). Salience of social issues in public discourse may determine whether or not issues expands on the government agenda (Koch-Baumgarten and Voltmer 2010). For example issue salience can determine voter turnout and choice preferences (Becker 1977). Our analysis suggests that print news media may have provided a method for establishing issue salience and coalescence, serving to bring the issue to the governmental agenda.

An indicator of policy image salience and coalescence is if the policy issue is being discussed in formal hearings (Baumgartner and Jones 2010). The first congressional hearing addressing IWPs was conducted in 1999 and addressed issues related to U.S. Department of Agriculture’s (USDA) policy for addressing wildlife transmission of diseases to domestic livestock, specifically brucellosis in IWPs (Senate 1999). This has been identified as a potentially significant issue facing agriculture and wildlife management (Miller et al. 2013, Bevins et al. 2014, Miller et al. 2017). However congressional hearings did not begin in earnest until 2005 and 2006 when ten hearings were held – over double from the previous five years. Hearings in these two years were largely related to potential animal agricultural impacts associated with classical swine fever, a swine disease with international trade implications for the U.S. swine industry (Paarlberg et al. 2009). These hearings coincided with a switch in the relative contribution of news media predictors and agriculture to policy activity. Indicating that print news media likely played an important role in forming the IWP policy image early in the policy process but interest groups played a more important role in formulating policies that would be implemented.

Once the issue of invasive IWPs was on the policy agenda and policy solutions were being developed, we found that the amount of agriculture in regions with IWPs was the most important predictor of the frequency of policy activity (Figure 4). The relative contribution to policy activity shifted from primarily media related to primarily agricultural after 2006 (see Figure S1.1 and S4). This indicates that agriculture may have been the primary driver influencing the development of potential policy solutions. Investigation of the policy documents revealed that there was an increased focus of IWP policy discourse on agricultural damage concerns indicating that agricultural interests had influence in both setting the policy agenda and the formation of policy solutions. While agriculture contributed relatively small amount to overall policy activity during the problem formation stage prior to policy image coalescence livestock agricultural contributed far greater than crop agriculture (see Figure 3 and panel B in Figure 4).

Previous studies have proposed that interest groups that are able to define the problem early in the issue emergence and problem formation stage tend to control future policy development even if new interest groups inter the policy arena (Schattschneider 1960, Baumgartner et al. 2009). Livestock agriculture involvement in policy formation may be driven by the potential for large economic losses – USD$5.8 billion - associated with a single livestock disease outbreak involving IWPs (foot and mouth disease) compared with the currently estimated USD$800 million in crop damage attributed to IWPs (Paarlberg et al. 2002, Pimentel et al. 2002, Pimental 2007, Anderson et al. 2016). Broadly this indicates that those interest groups with the greatest potential risk for damage had the greatest impact on the formation of IWP policy (Baumgartner et al. 2009). This effect may be even greater early in the emergence of an issue when fewer interest groups are involved and the ability for a single group to define the problem and the resulting policy is greater (Schattschneider 1960, Baumgartner et al. 2009).

Our approach may provide insights into the drivers of policy activity for wildlife and for invasive species such as feral horses or invasive fish, both of which receive significant media attention, are conflict-ridden, and impact wildlife and agriculture (Kincaid and Fletcher 2017, Kokotovich and Andow 2017). Using our approach for existing policy areas such as these may improve policy maker understanding of the drivers and importance of different interest groups overtime. This can then be used as a tool for improving stakeholder engagement or identifying interest groups that influence policy but have not been formally engaged by policy makers. A potentially more important application of our approach is to newly emerging policy areas in wildlife and invasive species. Early engagement by policy makers can be critical in defining successful policies before a policy area becomes grid locked. In addition, previous research indicates that interests groups whom engage early in a policy area often determine policy outcomes (Sapat 2004). Using our approach in combination with structured decision making techniques (Estévez et al. 2015) or frameworks that identifying stakeholder characteristics and synthesizing public preferences (Sharp et al. 2011) may improve proactive policy development by agencies avoiding policy development that is influenced by a single interest group.

This is the first analysis we are aware of that examines the relative contributions of media and institutional pressures on the development of invasive species policies at the Federal scale. In the case of IWPs, our model suggests that changes in co-occurrence of IWPs and agriculture over the last 29 years, likely resulting in increased problem severity (i.e. damage), contributed most to the eventual development of policy to mitigate the problems. This likely resulted from increasing industry pressure on agricultural agencies to protect or mitigate damage associated with IWPs. This was evidenced by both our model predictions and also the amount of consideration given to agricultural damage, specifically damage to livestock, caused by IWPs in congressional hearings and reports (GPO 2001, 2013). Livestock agricultural likely contributed more to the development of policy and has potential implications for implementation of programs to address IWP damage. Given the significant contribution of livestock agriculture to the formation and implementation of policy, national program objectives such as surveillance of IWPs for livestock diseases of concern and mitigation of risks associated with transmission of disease from IWPs to livestock are of particular importance and will require careful planning and implementation.

This study is based on a large search of government documents and news media data; therefore there are inherent constraints on inference. While our objective was to investigate the relative contribution of media and institutional pressures on national invasive species policy development, there are other potential drivers of policy activity. Previous studies have found that interest group access to congressional committees and advisory committees are influential in the development of policy (Balla and Wright 2001), although this is also influenced by the number of stakeholders in a policy area (Baumgartner et al. 2009). In our study we only considered three actors – livestock agriculture, crop agriculture, and the public – although there were likely additional actors that contributed to the generation of national policy such as conservation or sportsmen focused interest groups. While IWP policy appears to have gone through a continuum of policy development there is no standard quantitative approach for determining policy phases and investigating other policy processes may be of value. We did not consider policy processes such as policy diffusion (Berry and Berry 1999) or policy entrepreneurs (Mintrom and Norman 2009) that may have influenced national policy activity. These policy processes may also have contributed to the observed policy activity. While our study provides insights into drivers of policy activity addressing the invasive species interface, it could be enhanced by investigating these other mechanisms that may also be important in creation of policy. Additional extensions to our study could investigate the relative contribution of science (e.g. peer reviewed scientific papers) to policy activity compared with interest groups representing agriculture, conservation, and sportsmen. This may be of particular importance to better understand at which policy stage scientific findings have the greatest influence on policy development.

Given the scarcity of rigorous quantitative policy work for problems resulting from the interface of invasive species, wildlife, agriculture greater attention is needed to disentangle the mechanisms driving policy development. Although research has examined the influence of media on policy development (Baumgartner and Jones 2010), there remains a lack of information linking measures of public perception and institutional pressures specifically for the interface of wildlife, invasive species, and agricultural. Such information could provide valuable insight into the variability in policy approaches addressing these interactions. Analyses, such as the one we conducted, may provide improved understanding of which stakeholders have contributed most to policy activity and may be especially useful in better understanding complex policy systems. Additionally our approach may also provide early insight into emerging policy areas enabling proactive policy development by agencies or early engagement by scientists to find solutions before the policy area becomes grid locked. In addition linking the results of analyses such as ours with policy and program evaluation could provide a means of determining if the implemented policy and program match the original determinants of the policy. Policy makers can in turn use analyses such as these to better design policies that align with public interests and policy benefactors ensuring long term success of policies by incorporating all interests making programs more effective. (Loomis and Helfand 2001).

## Acknowledgements

We acknowledge the diligent efforts of research librarian Mary Foley who provided guidance on database searches. We also thank Drs. Robert Duffy, Sandra Davis, Dana Cole, Troy Bigelow, Dan Grear and Steve Sweeney for review and suggestions concerning hypotheses evaluated, analysis and discussion of findings. We also thank Dr. Terry Messmer and another anonymous reviewers for thoughtful suggestions that have improved this manuscript.

